# PIK-75 suppresses an SREBP-centred sterol programme in PIK3CA-altered small-cell lung cancer cells, but attribution to p110α remains unresolved

**DOI:** 10.64898/2026.10.08.757520

**Authors:** A Movaqar, A Yaghoubi, SAR Rezaee

## Abstract

**Background:** Class IA PI3K isoforms are candidate therapeutic targets in small-cell lung cancer (SCLC). The transcriptional consequences of isoform-directed inhibitors in SCLC cells have not been compared genome-wide.

**Methods:** We re-analysed GEO series GSE40564: PIK3CA-altered NCI-H69 cells treated for 24 h with vehicle, PIK-75 (p110α-directed) or TGX-221 (p110β-directed), three arrays per group (Affymetrix HG-U133 Plus 2.0). Differential expression used limma, with re-processing from raw files and spike-in-anchored normalisation as sensitivity analyses. Gene sets were assessed by effect size, competitive and rotation tests, and size-matched random-set nulls. Isoform dependency was examined in DepMap CRISPR screens. A PIK-75 response score was tested in 81 primary SCLCs against random-signature nulls.

**Results:** PIK-75 reduced SREBP-controlled sterol-and lipid-synthesis genes (SQLE, HMGCS1, HMGCR, LDLR, SCD; log2 fold-change −1.1 to −1.6). Cholesterol, mTORC1, hypoxia and glycolysis gene sets fell far more than size-matched random sets. E2F-target and homologous-repair genes fell more modestly, including BRCA1 (−0.91). The response was broad: 11,645 of 17,458 genes changed at FDR < 0.05, but only 316 decreased and 20 increased significantly beyond two-fold, an asymmetry that persisted under spike-in-anchored normalisation. TGX-221 changed 358 genes, none two-fold. Its set-level effects were four-to nine-fold smaller and depended on two of three replicates. PIK-75 did not induce canonical FOXO targets, and it inhibits kinases other than p110α, so the response cannot be attributed to p110α alone. In DepMap, SCLC lines depended more on PIK3CA than on PIK3CB (median Chronos −0.38 vs −0.15), as did other lineages. In patients, the response score was not associated with survival or subtype beyond what random gene signatures achieve (empirical p = 0.14 and 0.11).

**Conclusion:** PIK-75, but not TGX-221, suppresses an SREBP-centred anabolic transcriptional programme in NCI-H69 cells. Attributing this to p110α requires confirmation with selective inhibitors or genetic perturbation.

## 1. Introduction

Small-cell lung cancer (SCLC) is an aggressive neuroendocrine tumour with few targeted treatment options (1). Its genome is dominated by near-universal inactivation of TP53 and RB1, and recurrent activating alterations in druggable kinases are uncommon (2). Transcriptional subtypes defined by ASCL1, NEUROD1, POU2F3 and YAP1 have been proposed as a framework for therapeutic stratification (3), but pathway dependencies that could be exploited in the clinic remain poorly defined.

The class IA phosphoinositide 3-kinases (PI3Ks) are one candidate. Their catalytic isoforms are not interchangeable. p110α transmits most receptor-tyrosine-kinase and insulin signalling (4), whereas p110β becomes the dominant isoform chiefly in PTEN-deficient tumours (5, 6). Downstream, PI3K–AKT– mTORC1 signalling sustains anabolic transcription, including the SREBP-dependent synthesis of sterols and fatty acids (7, 8). Selective p110α inhibitors are approved for PIK3CA-mutant breast cancer (9). In SCLC, alterations of the PI3K pathway occur in a minority of tumours (2), and it is not established which isoform, if any, these tumours depend on.

Wojtalla et al. (10) addressed this question experimentally. They reported that p110α and p110β are overexpressed in primary SCLC, and that targeting p110α by RNA interference or with inhibitors impaired proliferation, survival and tumour growth, whereas targeting p110β was less effective. As part of that study, NCI-H69 cells were profiled on expression arrays after treatment with PIK-75, a p110α-directed inhibitor, or TGX-221, a p110β-directed inhibitor (11). The original report used these data mainly to describe changes in Bcl-2 family genes. The genome-wide responses to the two compounds have not been compared systematically.

Such a comparison needs care, for three reasons. First, PIK-75 is not selective: it inhibits DNA-PK (4) and many other kinases, including CDK9, a general regulator of transcription (12, 13). Second, NCI-H69 carries an in-frame PIK3CA deletion (p.Gly106_Arg108del) (14), a non-hotspot alteration that may influence baseline p110α pathway dependency. Third, when a treatment changes a large fraction of the transcriptome, standard normalisation and enrichment methods can create apparent effects that are not real.

We therefore re-analysed this dataset with three aims: to describe and compare the transcriptional responses to PIK-75 and TGX-221; to test how robust they are to data processing, normalisation and the choice of gene-set method; and to assess how far the PIK-75 response can be attributed to p110α. We then placed the findings in context using CRISPR dependency data from SCLC cell lines and expression data from primary SCLC. We show that PIK-75, but not TGX-221, suppresses an SREBP-centred anabolic programme, and we set out why this result cannot yet be assigned to p110α alone.

## 2. Methods

### 2.1 Dataset and model

We obtained GEO series GSE40564 (10, 15), generated on the Affymetrix Human Genome U133 Plus 2.0 array (GPL570; 54,675 probe sets). According to the GEO record, NCI-H69 SCLC cells (RRID:CVCL_1579) were treated for 24 h with DMSO, PIK-75 (100 nM) or TGX-221, three samples per group. The GEO record annotates TGX-221 at ’1 mM’. This is probably a typographical error for 1 µM, but we could not confirm the concentration from the record and therefore state no dose for TGX-221. The record does not state whether the three samples per group were independent experiments or parallel cultures.

NCI-H69 carries a heterozygous in-frame PIK3CA deletion (p.Gly106_Arg108del) according to Cellosaurus (16), which cites Shibata et al. (14). DepMap 24Q4 lists no PIK3CA hotspot call and no damaging PTEN mutation for this line. Sample-to-group assignment was taken from the GEO sample titles:

| Group | Treatment (primary target) | GEO samples |
| --- | --- | --- |
| DMSO | Vehicle | GSM996643, GSM996644, GSM996645 |
| PIK-75 | p110α-directed inhibitor, 100 nM | GSM996646, GSM996647, GSM996648 |
| TGX-221 | p110β-directed inhibitor | GSM996649, GSM996650, GSM996651 |

### 2.2 Array quality control and re-processing

Raw CEL files were inspected for scan date, probe-intensity distribution, RNA degradation slope, relative log expression and the Affymetrix hybridisation (bioB, bioC, bioD, cre) and polyA (lys, phe, thr, dap) spike-in controls. All nine arrays were scanned on one day within 2 h 14 min, in group order.

The main analysis used the submitter-processed RMA values (17). Two re-analyses from CEL files (affy package) (18) tested their robustness. The first was RMA: convolution background correction, quantile normalisation and median polish. The second was spike-in-anchored normalisation: the same pipeline without the quantile step, with each array shifted so that the mean log2 signal of the Affymetrix hybridisation (bioB, bioC, bioD, cre) and polyA (lys, phe, thr, dap) control probe sets was equal across arrays. This relaxes the assumption that most genes are unchanged. It is a sensitivity analysis only. In the standard Affymetrix protocol the polyA controls are added to the RNA sample before labelling and the hybridisation controls to the hybridisation mixture, so they reference signal to the amount of material processed, not to cell number, and the result should not be read as absolute transcript abundance per cell. The GEO record does not describe how the controls were added; both sets were detected at near-constant intensity on all nine arrays (Table S11).

### 2.3 Probe filtering and gene-level collapsing

We removed control probes (AFFX-), unannotated probes and probes mapped to more than one gene symbol. We kept probes whose intensity exceeded the 25th percentile of all values in at least three samples (32,709 probes). Each gene was represented by its probe with the highest mean intensity across all samples (“MaxMean” (19); 17,458 genes). The rule was fixed before any contrast was computed.

As a sensitivity analysis, probes were averaged per gene (limma::avereps). For PIK-75 vs DMSO, log2 fold-changes agreed between rules (Pearson r = 0.91, Spearman ρ = 0.92). Among genes significant under both rules, 98.5% had the same sign. The sets of two-fold differentially expressed genes overlapped less (781 vs 485 genes; Jaccard 0.51), because averaging dilutes fold-changes (Table S6). Genes whose direction depends on the probe are flagged in the text.

### 2.4 Differential expression

We fitted a means model (∼0 + group) with limma (20) and extracted three contrasts:

| Contrast | Definition | Question answered |
| --- | --- | --- |
| C1 | PIK-75 – DMSO | Transcriptional effect of PIK-75 |
| C2 | TGX-221 – DMSO | Transcriptional effect of TGX-221 |
| C3 | TGX-221 – PIK-75 | Difference between the two responses (C2 – C1); a derived contrast that adds no independent degree of freedom |

Moderated t-statistics used empirical Bayes shrinkage with an intensity trend and robust hyperparameter estimation (21, 22). P-values were adjusted by the Benjamini–Hochberg method (23). We report genes at FDR < 0.05, and genes significantly beyond a fold-change threshold using limma::treat (24) at 1.5-fold and two-fold. Counts at FDR < 0.05 with |log2FC| ≥ 1 are given for comparison.

For C2, we assessed dependence on individual arrays in two ways: empirical array weights (25), and leaving out each TGX-221 replicate in turn.

### 2.5 Exploratory analysis and visualisation

Principal component analysis used the 1,000 most variable genes, centred but not scaled. The heatmap shows row z-scores of the 60 genes with the largest moderated F-statistic across C1 and C2 (FDR < 0.05 and |log2FC| ≥ 1 in at least one contrast), for all nine samples.

### 2.6 Gene-set analysis

Four complementary measures, because PIK-75 changes most of the transcriptome.

#### Effect size

The mean log2 fold-change of all genes in a set, and the percentage of its genes that decreased.

#### Gene-set tests

Self-contained rotation tests (limma::mroast, 4,999 rotations) (26) and competitive tests adjusted for inter-gene correlation (limma::camera) (27) were applied to the 15 gene sets listed in Table S16. Self-contained tests saturate when most genes change, so rotation-test results are reported for completeness and are not used alone to support a claim.

#### Random-set null

The mean log2 fold-change of each set was compared with the means of 10,000 random gene sets of the same size, drawn without replacement from the 17,458 tested genes. We report a z-score (observed mean minus the mean of the random sets, divided by their standard deviation) and a two-sided empirical p-value (Table S23).

#### Competitive enrichment

Pre-ranked GSEA (fgsea) (28, 29) on moderated t-statistics, using MSigDB Hallmark (30) and Reactome (31) sets of 15–500 genes. A positive normalised enrichment score (NES) means that a set decreased less than the rest of the transcriptome. It does not establish induction.

### 2.7 Pathway and transcription-factor activity

Pathway activity was inferred from PROGENy footprints (32) (top 500 genes per pathway; multivariate linear model in decoupleR)(33). Transcription-factor activity was inferred from DoRothEA regulons of confidence A–C (34) (univariate linear model). PROGENy scores were compared with a null distribution from 200 permutations of gene labels. Because regulons overlap the genes examined individually, we counted how many of the SREBP targets in Table 2 belong to the SREBF1 and SREBF2 regulons.

### 2.8 mRNA half-life and LINCS signature analyses

To test whether the PIK-75 response resembles a general block of transcription, we correlated log2 fold-changes with consensus human mRNA half-lives (35) (12,341 genes matched). A block of transcription predicts that short-lived mRNAs fall most.

We also stratified genes by half-life. The shortest-lived 5% of transcripts by consensus half-life score (the score has no time units) were compared with all other transcripts, for log2 fold-change (Mann–Whitney test) and for the percentage reduced at least two-fold at FDR < 0.05 (Table S15b). In addition, the PIK-75 response was compared with 5,425 compound signatures from LINCS L1000 (36) (consensus signature library from SigCom LINCS, obtained through Enrichr (37, 38) For each compound, similarity was the mean moderated t-statistic of the genes in its ‘up’ signature minus that of the genes in its ‘down’ signature (at least 20 matched genes each), standardised to a z-score across compounds and expressed as a percentile; high values indicate a similar response. We examined transcription and CDK9 inhibitors (dactinomycin, dinaciclib, alvocidib, AT-7519) (39) mTOR kinase and PI3K inhibitors (Torin-1, Torin-2, AZD-8055, alpelisib) (40–42), the DNA-PK inhibitor NU-7441 and the LINCS signature of PIK-75 itself (Table S22).

### 2.9 DepMap dependency analysis

CRISPR gene-effect (Chronos) (43) scores, model annotations and mutation matrices came from DepMap Public 24Q4 (44, 45). SCLC lines were those with OncotreeCode SCLC and CRISPR data (n = 25). NSCLC lines had OncotreeCode LUAD, LUSC or NSCLC (n = 80). A line was called dependent at Chronos < −0.5. Differences are reported as Hodges–Lehmann shifts with 95% confidence intervals. The within-SCLC comparison was repeated after excluding lines with a PIK3CA hotspot or a damaging PTEN mutation. NCI-H69 was not screened.

### 2.10 Primary SCLC cohort

RNA-seq and clinical data for the UCologne cohort (2) (n = 81) were retrieved through the cBioPortal API (46, 47) and log2(x + 1)-transformed. A PIK-75 response score was computed per tumour (Table S10): the mean z-score of the genes decreased by PIK-75 minus the mean z-score of the increased genes (C1; FDR < 0.05 and |log2FC| ≥ 1).

Subtype was assigned by the highest z-scored marker among ASCL1, NEUROD1, POU2F3 and YAP1 (3). A neuroendocrine score was computed from the 50-gene panel of Zhang et al. (48) as the mean z of neuroendocrine genes minus the mean z of non-neuroendocrine genes. Overall survival was modelled by Cox regression per SD of the score, adjusted for stage (I–II vs III–IV), and additionally for neuroendocrine score, age and sex.

Specificity was assessed against 1,000 random gene signatures of the same size, drawn from genes expressed in NCI-H69, and against 1,000 signatures matched for correlation with the neuroendocrine score (49). The empirical p-value is the fraction of random signatures with a test statistic at least as extreme as the observed one.

### 2.11 Software, data and code availability

Analyses used R 4.4.3 (50) with limma 3.62.1, affy 1.84.0, GEOquery 2.74.0 (51), fgsea 1.32.2, msigdbr 26.1.1, progeny 1.28.0, dorothea 1.18.0, decoupleR 2.12.0, survival 3.8.12 (52), data.table 1.18.6.1, ggplot2 4.0.3 and pheatmap 1.0.13. The random seed was 20260930. The random-set null, the stratified half-life test and the LINCS comparison were run in Python 3.11.15 with pandas 3.0.2, NumPy 2.4.4 and SciPy 1.17.1 (seed 20261005). The analysis scripts, all intermediate tables and a full sessionInfo() are available at https://github.com/Aref-Movaqar/sclc-pi3k-transcriptomics, archived at Zenodo https://doi.org/10.5281/zenodo.23154076.

**Table 1.** Differential expression by contrast (17,458 genes tested).

| Contrast | FDR < 0.05 | FDR < 0.05 and ≥ 2-fold (up / down) | Significantly > 2-fold by treat (up / down) | Spike-in-anchored, ≥ 2-fold (up / down) |
| --- | --- | --- | --- | --- |
| C1: PIK-75 vs DMSO | 11,645 | 51 / 730 | 20 / 316 | 21 / 1,504 |
| C2: TGX-221 vs DMSO | 358 | 0 / 0 | 0 / 0 | 0 / 0 |

## 3. Results

### 3.1 PIK-75 and TGX-221 produce very different transcriptomic responses

Array quality was adequate. Hybridisation and polyA spike-in controls were constant across the nine arrays (log2 11.2–11.4 and 7.9–8.1), and median relative log expression was within 0.04 of zero (Table S11). One PIK-75 array (PIK75_5) showed more 5′ signal loss for ACTB than the others. Re-processing from CEL files reproduced the submitter values (r = 1.000 for log2 fold-changes; Table S12).

PIK-75 dominated the variation. The first principal component explained 96.2% of variance and separated the PIK-75 replicates from all other samples (Figure 2A). DMSO and TGX-221 samples intermixed, and one TGX-221 replicate clustered with the controls (Figure 1).

**Figure 1.**
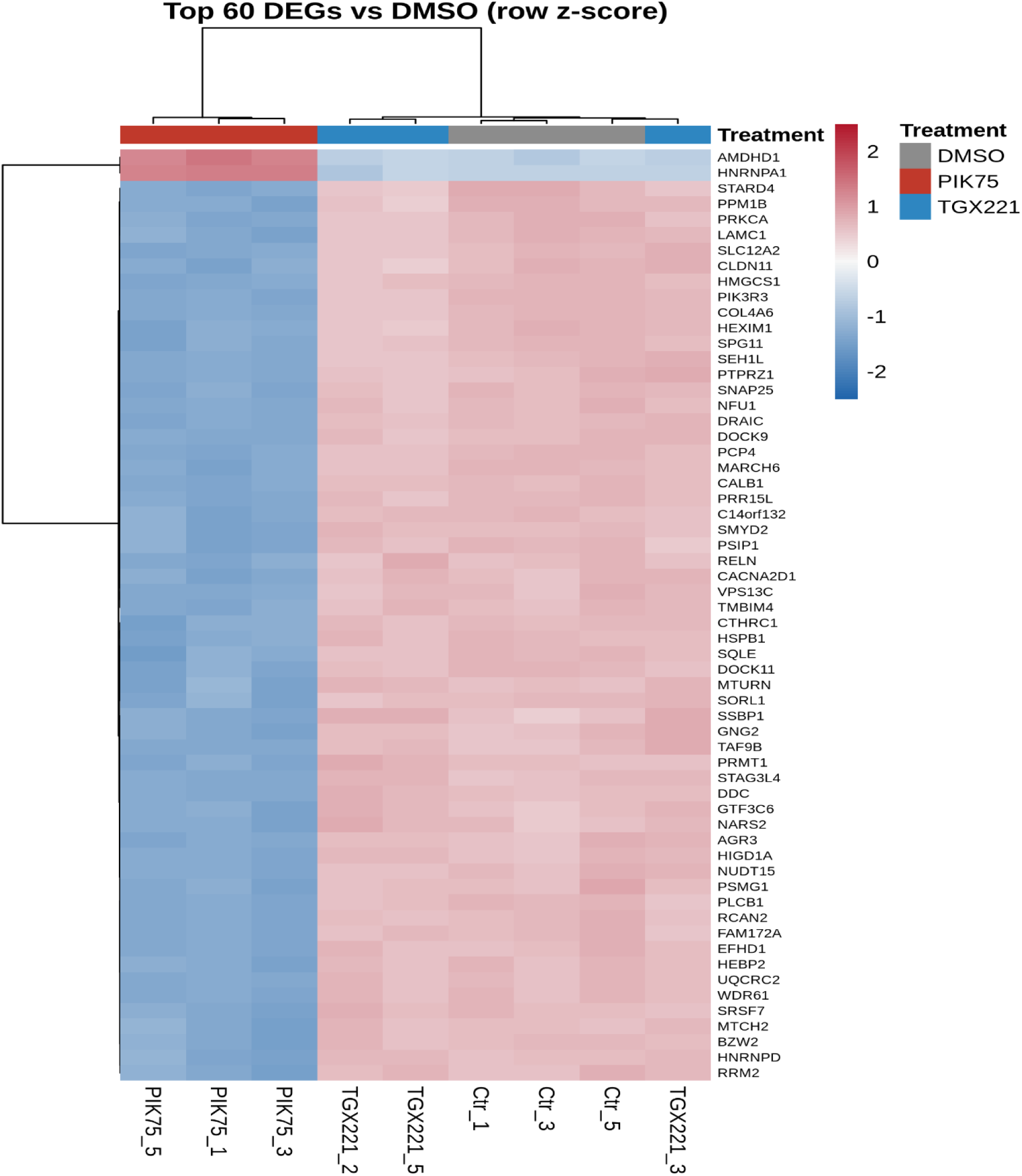
PIK-75 drives a coherent transcriptional response that TGX-221 does not reproduce. Row z-scored expression of the 60 genes with the largest moderated F-statistic across the PIK-75 vs DMSO and TGX-221 vs DMSO contrasts (FDR < 0.05 and |log2FC| ≥ 1 in at least one contrast), shown for all nine samples. Rows are clustered by correlation distance and columns by Euclidean distance. The annotation bar gives treatment. TGX-221 replicates are not separated from DMSO; replicate TGX221_3 clusters with the controls.

**Figure 2.**
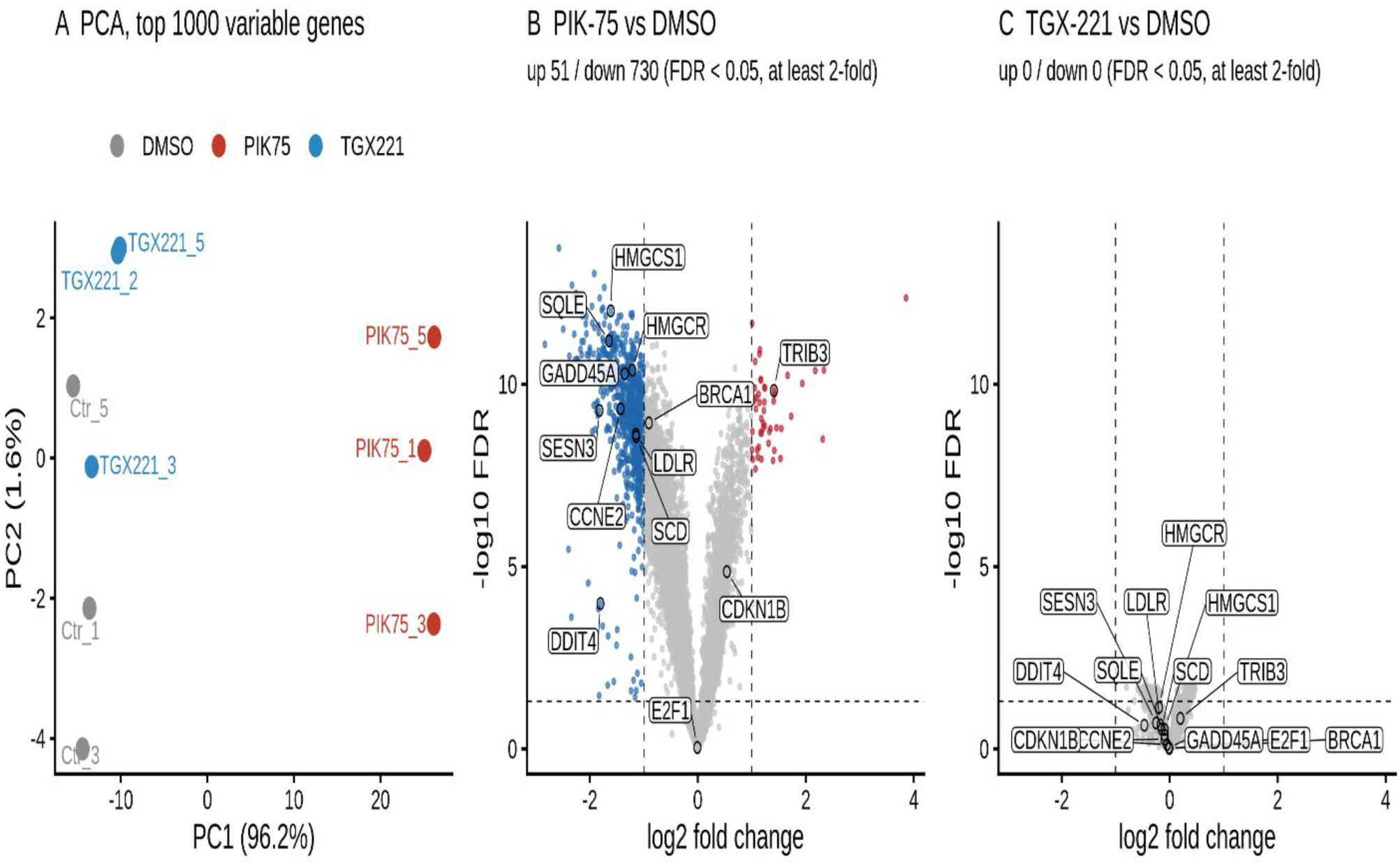
PIK-75 dominates sample variance. (A) Principal component analysis of the 1,000 most variable genes. (B, C) Volcano plots for PIK-75 vs DMSO and TGX-221 vs DMSO on identical axes. Dashed lines mark FDR = 0.05 and |log2FC| = 1. Red and blue points are genes up-or down-regulated beyond both thresholds, with counts in each panel title. Selected genes discussed in the text are labelled.

PIK-75 changed two-thirds of expressed genes at FDR < 0.05 (Table 1; Table S1; Figure 2B). With three arrays per group and a shift this broad, that count reflects statistical power and normalisation as much as biology (it was 8,009 with spike-in anchoring; Table S12), so interpretation below rests on effect sizes and fold-change-thresholded tests. Large changes were mostly decreases: 316 genes fell and 20 rose significantly beyond two-fold (Table S13). This asymmetry did not depend on quantile normalisation. With spike-in-anchored normalisation there were 1,504 two-fold decreases and 21 increases, and gene ranking was preserved (r = 0.975 with the main analysis). Across all genes the response was not one-sided: 40% of log2 fold-changes were negative in the main analysis and 57% with spike-in anchoring. TGX-221 changed 358 genes at FDR < 0.05 and none two-fold under any method (Figure 2C).

### 3.2 PIK-75 suppresses SREBP-dependent sterol synthesis and an mTORC1–hypoxia– glycolysis programme

Sterol-and lipid-synthesis genes were among the most strongly reduced transcripts (Table 2). Seven fell more than two-fold (SQLE, HMGCS1, HMGCR, LDLR, SCD, LSS and FDFT1; log2FC −1.0 to −1.6; all FDR < 0.001). To avoid relying on genes chosen after seeing the data, we also examined the Reactome set ‘Activation of gene expression by SREBF (SREBP)’ (R-HSA-2426168; 41 genes tested): 25 decreased and 5 increased at FDR < 0.05 (mean log2FC −0.37 vs −0.06 with TGX-221). Seven of the ten SREBP targets in Table 2 belong to this set; LDLR, LSS and INSIG1 do not.

**Table 2.** Selected genes (log2 fold-change vs DMSO). The probe is the highest-intensity probe per gene. All PIK-75 values with |log2FC| ≥ 0.3 have FDR < 0.001. No TGX-221 value is significant. Further gene panels are in Table S17.

| Group | Gene | PIK-75 | PIK-75, spike-in-anchored | TGX-221 |
| --- | --- | --- | --- | --- |
| SREBP targets | SQLE | -1.64 | -1.96 | -0.09 |
| SREBP targets | HMGCS1 | -1.61 | -2.05 | -0.09 |
| SREBP targets | HMGCR | -1.21 | -1.55 | -0.20 |
| SREBP targets | LDLR | -1.15 | -1.44 | -0.17 |
| SREBP targets | SCD | -1.14 | -1.52 | -0.15 |
| SREBP targets | LSS | -1.04 | -1.21 | -0.23 |
| SREBP targets | FDFT1 | -1.01 | -1.39 | -0.14 |
| SREBP targets | ACACA | -0.83 | -1.07 | -0.10 |
| SREBP targets | INSIG1 | -0.82 | -1.27 | -0.01 |
| SREBP targets | FASN | -0.30 | -0.55 | -0.15 |
| Cell cycle / repair | CCNE2 | -1.43 | -1.74 | -0.10 |
| Cell cycle / repair | BRCA1 | -0.91 | -1.19 | -0.01 |
| Cell cycle / repair | BRCA2 | -0.42 | -0.51 | -0.19 |
| Cell cycle / repair | RAD51 | -0.33 | -0.60 | -0.06 |
| Cell cycle / repair | E2F1 | -0.01 | -0.25 | -0.01 |
| Cell cycle / repair | MKI67 | +0.11 | -0.24 | -0.06 |
| FOXO targets | SESN3 | -1.82 | -2.10 | -0.25 |
| FOXO targets | GADD45A | -1.35 | -1.62 | -0.01 |
| FOXO targets | CITED2 | -1.32 | -1.54 | -0.32 |
| FOXO targets | CDKN1B | +0.54 | +0.10 | -0.05 |
| ATF4 targets | TRIB3 | +1.41 | +1.10 | +0.20 |
| ATF4 targets | SLC7A5 | +0.73 | +0.36 | -0.03 |
| ATF4 targets | SESN2 | +0.62 | +0.52 | +0.11 |
| ATF4 targets | DDIT4 | -1.80 | -2.17 | -0.47 |

None of the ten SREBP targets in Table 2 changed by more than 0.23 log2 units with TGX-221. The decreases were larger still with spike-in-anchored normalisation. SREBF2 mRNA itself was unchanged (−0.03), and SREBF1 fell slightly (−0.32).

At the gene-set level, Reactome cholesterol biosynthesis showed the largest effect (mean log2FC −0.64; 88% of genes decreased), followed by Hallmark cholesterol homeostasis (−0.38), mTORC1 signalling (−0.33), hypoxia (−0.30) and glycolysis (−0.28; Table 3; Tables S2, S3 and S16; Figure 3). Because widespread repression can saturate self-contained tests, we evaluated mean effect sizes against empirical null distributions from 10,000 size-matched random gene sets (Table S23). All primary metabolic programmes fell far beyond random expectation (cholesterol biosynthesis: z = −6.6; cholesterol homeostasis: z = −6.4; mTORC1: z = −8.9; hypoxia: z = −8.0; glycolysis: z = −7.2; all empirical p = 0.0001). Conversely, sets that appeared nominally altered under competitive or self-contained tests failed against random nulls (unfolded protein response: z = −0.3, p = 0.80; DNA repair: z = −1.5, p = 0.12; ATF4 stress response: z = −0.9, p = 0.37). Inferred PI3K pathway activity fell sharply (PROGENy t = −7.57, permutation p < 0.01; Figure 4A; Tables S4 and S18), and inferred SREBF1 and SREBF2 activity decreased in tandem (t = −6.81 and −6.09; Figure 4B; Table S5). These regulon scores are not independent of the gene-level result: the SREBF1 regulon contains nine of the ten SREBP targets in Table 2, and the SREBF2 regulon five.

**Table 3.** Gene-set effect sizes. Mean log2 fold-change of all genes in the set; values in parentheses are z-scores against 10,000 size-matched random gene sets (Table S23). The transcriptome-wide mean is −0.05 for PIK-75 and 0.00 for TGX-221. The rotation-test columns give the FDR across the 15 sets tested; these tests saturate under a global shift and are shown for completeness. Sets are from the MSigDB Hallmark collection unless marked Reactome.

| Gene set (n genes) | PIK-75 mean log2FC (z) | % of genes decreased, PIK-75 | Rotation test, PIK-75 | TGX-221 mean log2FC (z) | Rotation test, TGX-221 | NES, PIK-75 / TGX-221 |
| --- | --- | --- | --- | --- | --- | --- |
| Cholesterol biosynthesis, Reactome (25) | −0.64 (−6.6) | 88 | 0.0002 | −0.07 (−2.5) | 0.15 | −2.16 / −1.58 |
| Cholesterol homeostasis (70) | −0.38 (−6.4) | 69 | 0.0002 | −0.04 (−2.8) | 0.064 | −1.94 / −1.46 |
| mTORC1 signalling (188) | −0.33 (−8.9) | 66 | 0.0002 | −0.06 (−6.1) | 0.15 | −1.83 / −1.96 |
| Hypoxia (186) | −0.30 (−8.0) | 58 | 0.0002 | −0.08 (−8.7) | 0.15 | −1.72 / −2.13 |
| Glycolysis (189) | −0.28 (−7.2) | 60 | 0.0002 | −0.06 (−6.0) | 0.15 | −1.74 / −1.78 |
| Homology-directed repair, Reactome (95) | −0.24 (−4.2) | 66 | 0.0002 | −0.04 (−3.1) | 0.30 | −1.58 / −1.57 |
| E2F targets (191) | −0.18 (−4.4) | 61 | 0.0007 | −0.02 (−2.2) | 0.57 | −1.49 / −1.16 |
| G2M checkpoint (185) | −0.14 (−2.9) | 57 | 0.0005 | −0.03 (−2.7) | 0.30 | −1.35 / −1.37 |
| DNA repair (141) | −0.10 (−1.5) | 48 | 0.056 | −0.02 (−1.1) | 0.65 | −1.17 / −0.86 |
| Unfolded protein response (100) | −0.06 (−0.3) | 56 | 0.26 | −0.05 (−3.6) | 0.17 | +1.51 / −1.38 |
| Translation initiation, Reactome (97) | +0.08 (+2.9) | 23 | 0.0005 (up) | +0.02 (+2.2) | 0.15 | +2.64 / +2.05 |

**Figure 3.**
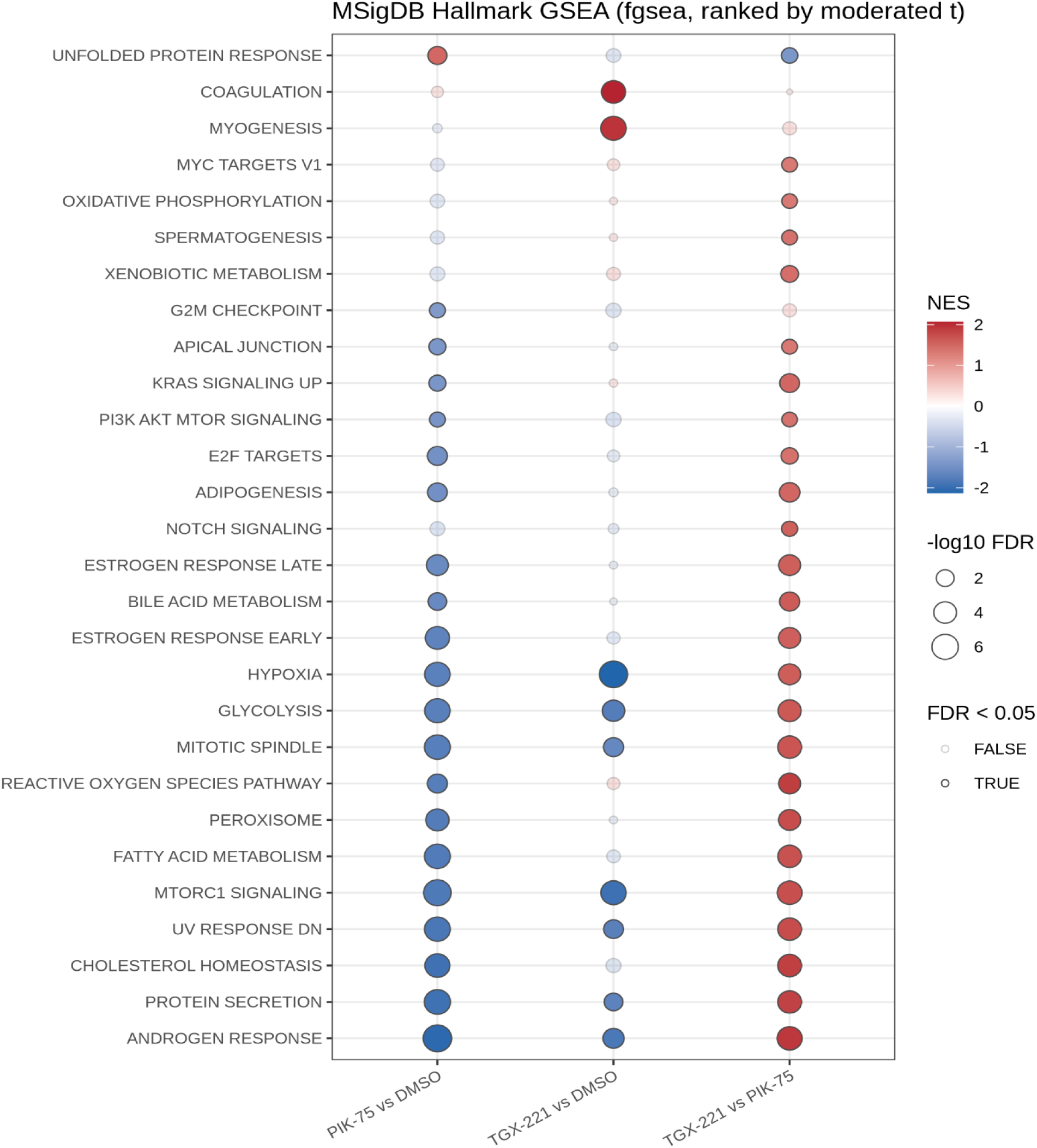
Hallmark gene-set enrichment for the three contrasts. Pre-ranked GSEA (fgsea) on moderated t-statistics for all 17,458 genes. The figure shows every Hallmark set with FDR < 0.05 in at least one contrast. Fill gives the normalised enrichment score (NES) and point size −log10 FDR. Faded points are not significant (FDR ≥ 0.05). Sets are ordered by NES in the PIK-75 vs DMSO contrast. NES measures enrichment relative to the rest of the transcriptome, not effect size: a positive NES can occur for a set that decreased less than average. Effect sizes are in Table 3.

**Figure 4.**
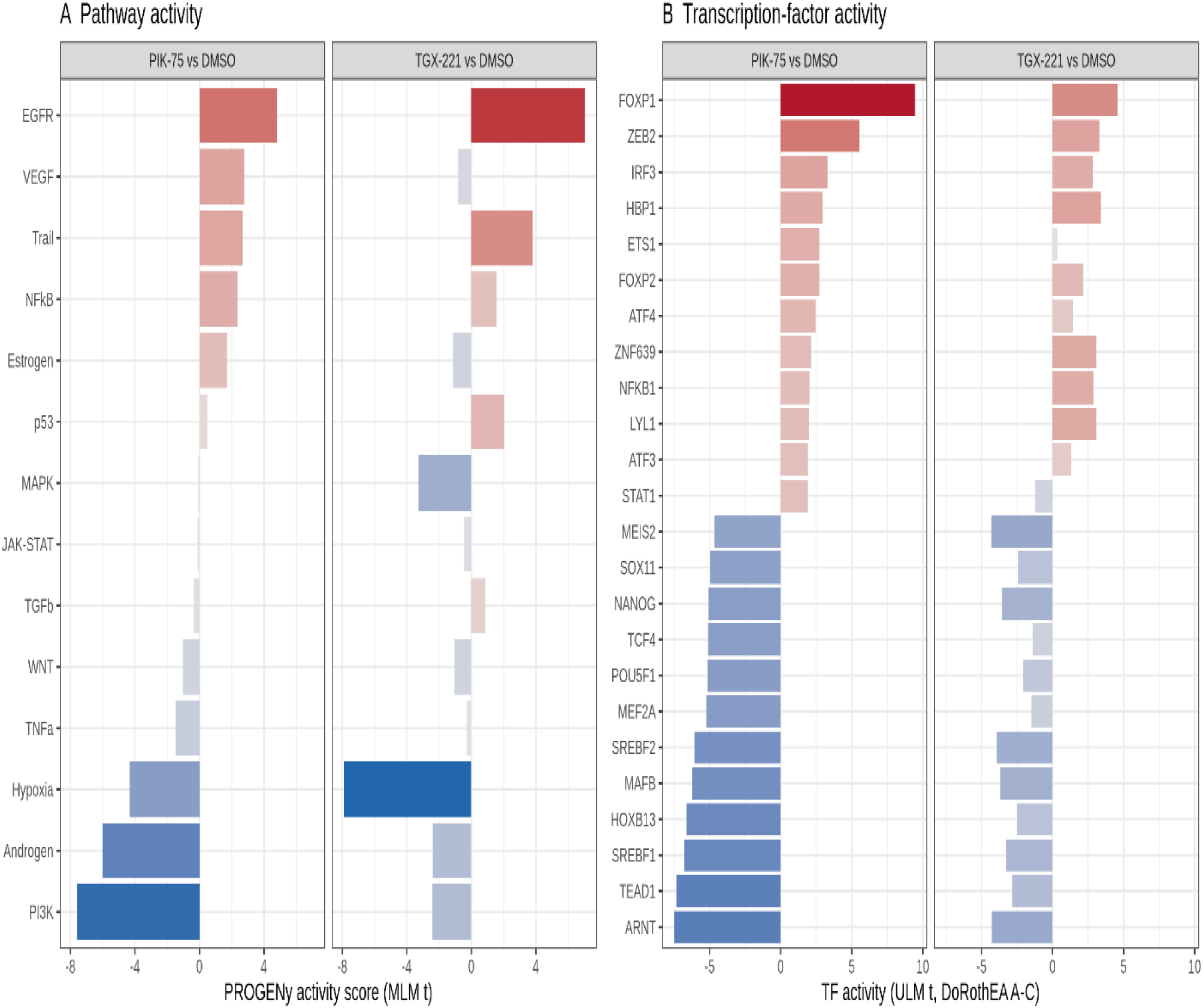
Inferred pathway and transcription-factor activity. (A) PROGENy pathway activity (multivariate linear model on the top 500 footprint genes per pathway) for PIK-75 vs DMSO and TGX-221 vs DMSO. (B) DoRothEA (confidence A–C) transcription-factor activity by univariate linear model: the 12 most decreased and 12 most increased regulators after PIK-75, with the TGX-221 values shown for comparison. Scores are inferred from target-gene changes and are not direct measurements.

E2F-target and homologous-repair gene sets decreased less (mean log2FC −0.18 and −0.24). BRCA1 fell by 0.91 log2 units, concordantly on both probes, CCNE2 by 1.43, and BRCA2 and RAD51 by 0.33–0.42. E2F1, MKI67, PCNA and TOP2A were unchanged, so the data do not show a general proliferative arrest. The Hallmark DNA Repair set was not significant (rotation-test FDR 0.056).

### 3.3 Evidence for and against an on-target p110α mechanism

Three observations fit inhibition of PI3K signalling. The PI3K footprint score fell. The mTORC1-and SREBP-dependent programmes that PI3K–AKT signalling is known to sustain decreased. ERK-pathway output genes rose modestly (DUSP6 +0.56, ETV4 +0.37), as expected when PI3K inhibition relieves feedback on receptor tyrosine kinases (53, 54).

Three observations do not. First, canonical FOXO targets, which AKT inhibition should induce (55), decreased: SESN3, GADD45A and CITED2 each fell more than two-fold, and inferred FOXO3 activity fell (t = −2.90). Only CDKN1B rose (+0.54). Second, the EGFR footprint score increased with both compounds (t = +4.82 for PIK-75 and +7.03 for TGX-221), so it is not specific to PIK-75. Third, the response spanned two-thirds of expressed genes, which is broader than expected from one signalling pathway.

We tested whether the response resembles a general block of transcription, as off-target inhibition of CDK9 would produce. The half-life analysis does not support a simple global shutdown. Log2 fold-change correlated only weakly with mRNA half-life (Spearman ρ = −0.065; Table S15), and the shortest-lived 5% of mRNAs were less reduced than the rest: 2.8% fell at least two-fold, compared with 5.3% of all other transcripts (Mann–Whitney p = 1.3 × 10⁻⁵ for log2 fold-change; Table S15b). This test is weak at 24 h and does not exclude a contribution from CDK9 inhibition.

The LINCS comparison was not decisive (Table S22). The PIK-75 response was dissimilar to most transcription and CDK9 inhibitor signatures (dactinomycin: percentile 0.3, z = −2.86; dinaciclib: percentile 1.3, z = −2.24; alvocidib: percentile 10.3, z = −1.23), although not to the multi-CDK inhibitor AT-7519 (percentile 91.6, z = +1.36). The DNA-PK inhibitor NU-7441 was the least similar of all 5,425 compounds (z = −4.25). The response was similar to ATP-competitive mTOR kinase inhibitors (Torin-2: percentile 99.5, z = +2.80; Torin-1: percentile 98.6, z = +2.22; AZD-8055: percentile 96.2, z = +1.76). However, it was also dissimilar to the selective p110α inhibitor alpelisib (percentile 6.5, z = −1.47) and to the LINCS signature of PIK-75 itself (percentile 7.9, z = −1.38). A comparison that does not recover the compound’s own signature cannot identify its mechanism: the LINCS consensus signatures are aggregated across cell lines, doses and exposure times and rest on 978 directly measured genes. We therefore read the result as consistent with a transcriptional state resembling mTOR kinase inhibition, not as evidence of mechanism.

Apparent increases need care. The unfolded-protein-response set had a positive NES but a mean log2 fold-change of −0.06, and it was not significant in the rotation test. The translation-initiation and GCN2 sets are 68–80% ribosomal-protein genes, which rose by less than 0.1 log2 units and fell under spike-in anchoring. A few ATF4 targets did rise, most clearly TRIB3 (+1.41, on both probes). The Reactome ATF4-target set as a whole decreased (mean −0.13). We therefore do not claim an integrated stress response.

### 3.4 TGX-221 produces a small, replicate-dependent response

TGX-221 changed no gene two-fold. Set-level shifts pointed the same way as with PIK-75 but were four-to nine-fold smaller in effect size (Table 3). No set reached significance in rotation tests (lowest FDR 0.064). The mTORC1, hypoxia and glycolysis shifts were significant under camera (FDR 0.02–0.05) and exceeded random-set expectation (z = −6.0 to −8.7; Table 3), so the shift in the full data is small but not random. Whether it reflects a drug effect is uncertain, because it depended on individual replicates (see below). The similar NES for mTORC1, hypoxia and glycolysis in the two contrasts reflect ranking within each contrast, not similar magnitude.

The response rested on two of three replicates. Removing TGX221_2 or TGX221_5 left no significant genes. Removing TGX221_3, which clustered with the controls, raised the count to 970, of which 351 were among the original 358 (Table S14). With empirical array weights, 1,961 genes were significant. The data therefore cannot distinguish a weak uniform effect from variable drug exposure across replicates. The original study reported that targeting p110β affected SCLC cell proliferation less than targeting p110α (10).

### 3.5 Dependency and clinical context

#### DepMap

Across 25 SCLC lines, PIK3CA knockout reduced fitness more than PIK3CB knockout (median Chronos −0.38 vs −0.15; paired shift −0.21, 95% CI −0.30 to −0.11; Figure 5A; Table 4; Table S8). The hierarchy held after excluding the one line with a PIK3CA hotspot mutation and the four with damaging PTEN mutations (n = 20; −0.42 vs −0.16; p = 2 × 10⁻⁶; Table S19). The four PTEN-mutant lines showed the lowest PIK3CB scores (median −0.29). PIK3CA dependency did not differ between SCLC and NSCLC lines (shift 0.04, 95% CI −0.05 to 0.14), so the hierarchy is not specific to SCLC. Genetic dependency is also not the same as sensitivity to an inhibitor.

**Figure 5.**
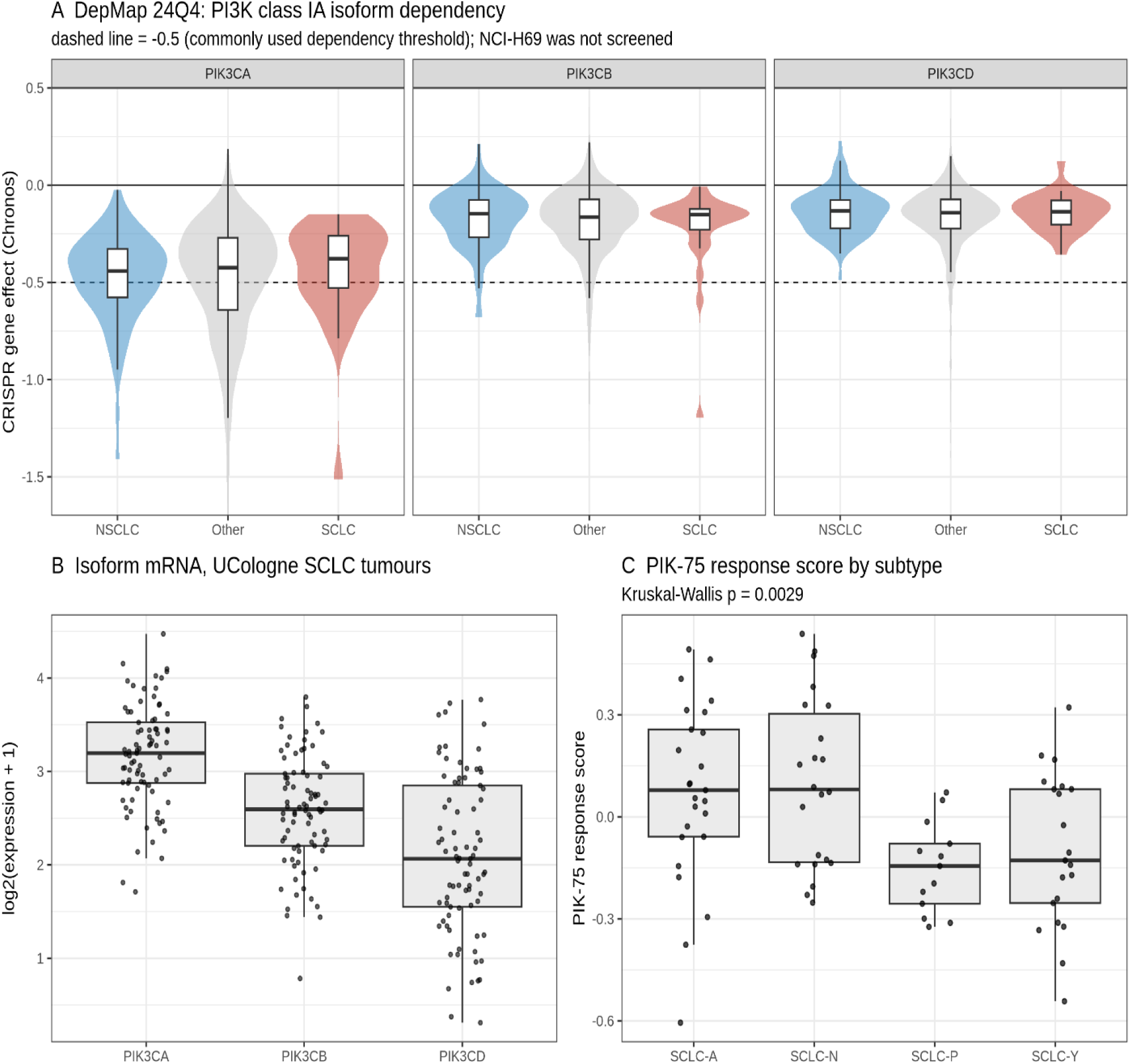
Dependency and clinical context. (A) DepMap 24Q4 CRISPR gene effect (Chronos) for PIK3CA, PIK3CB and PIK3CD in SCLC (n = 25), NSCLC (n = 80) and other (n = 1,073) cell lines. The dashed line marks −0.5. The y-axis is truncated at −1.6. (B) PIK3CA, PIK3CB and PIK3CD mRNA in UCologne primary SCLC (n = 81). (C) The PIK-75 response score (mean z of PIK-75-repressed genes minus mean z of PIK-75-induced genes) by dominant-marker subtype. Kruskal–Wallis p = 0.0029, which is not more extreme than random signatures of the same size achieve (empirical p = 0.11).

**Table 4.** DepMap 24Q4 Chronos gene effect.

| Gene | SCLC (n = 25) median / % dependent | NSCLC (n = 80) median / % dependent | Other (n = 1,073) median / % dependent |
| --- | --- | --- | --- |
| PIK3CA | $-0.38$ / 40% | $-0.44$ / 40% | $-0.42$ / 39% |
| PIK3CB | $-0.15$ / 8% | $-0.15$ / 4% | $-0.16$ / 6% |
| PIK3CD | $-0.14$ / 0% | $-0.13$ / 0% | $-0.14$ / 4% |

#### Primary SCLC (UCologne, n = 81)

The PIK-75 response score was not specifically associated with outcome or subtype. In a stage-adjusted Cox model the hazard ratio was 1.36 per SD (95% CI 1.02–1.80; 77 patients, 48 deaths; Table S9). However, 16% of random signatures of the same size reached nominal significance in this cohort, and the observed association was not unusual among them (empirical p = 0.14; Table S21). The same held for the difference between transcriptional subtypes (Kruskal–Wallis p = 0.003; empirical p = 0.11; Figure 5C).

The score was not a linear proxy for neuroendocrine differentiation across the full cohort (overall Spearman ρ = 0.09 with the 50-gene neuroendocrine score), and adjusting for that score left the hazard ratio unchanged (1.37; 1.03–1.84; Table S20). While the score differed across subtypes (higher in SCLC-A and -N than in -P and -Y; Kruskal–Wallis p = 0.003), within each individual subtype the score was inversely correlated with the neuroendocrine score (ρ = −0.49 in SCLC-A, −0.24 in SCLC-N, and −0.31 in SCLC-Y). This intra-subtype negative relationship cancels the between-subtype elevation, explaining the near-zero global correlation.

Six tumours with expression data carried mutations in PIK3CA, PTEN, AKT2 or PIK3R1; their scores did not differ from the rest (p = 0.84). PIK3CA mRNA exceeded PIK3CB mRNA (median log2 3.20 vs 2.60; Figure 5B). The score did not correlate more strongly with PIK3CA than with PIK3CB (difference in ρ 0.11; 95% CI −0.09 to 0.32). We conclude that the cohort provides no independent support for the clinical relevance of the PIK-75 response.

## 4. Discussion

### 4.1 Summary

In NCI-H69 cells, PIK-75 had a far larger transcriptional effect than TGX-221. Its clearest component was the loss of SREBP-dependent sterol-and lipid-synthesis genes, together with mTORC1-, hypoxia-and glycolysis-associated transcripts. E2F and homologous-repair genes decreased more modestly. TGX-221 produced small shifts in the same direction that depended on individual replicates. These are differences between two compounds at one dose in one cell line. They do not by themselves establish a difference between p110α and p110β.

We therefore separate three levels of evidence. Observed: PIK-75 produces a broad transcriptional response in NCI-H69 cells, whose most coherent component is reduced expression of SREBP target genes. Consistent with mechanism: this component is what reduced PI3K–AKT–mTORC1 signalling would be expected to produce. Not established: that the response is caused by inhibition of p110α.

### 4.2 What can be attributed to p110α

Several features of the experiment limit attribution to p110α:

#### Selectivity of PIK-75

At 200 nM, PIK-75 inhibited 27 kinases in addition to p110α in a 451-kinase screen, including CDK7 and CDK9 (13). It also inhibits DNA-PK (4) and suppresses CDK9-dependent transcription in leukaemia cells (12). The dataset used 100 nM.

#### Breadth and direction of the response

PIK-75 changed two-thirds of expressed genes, and 94% of the changes significantly beyond two-fold were decreases. Canonical FOXO targets fell, although AKT inhibition should induce them.

#### Unequal comparison

The two compounds differ in potency and selectivity, and were each used at a single dose. In the original study, targeting p110β affected SCLC cell proliferation less than targeting p110α (10), Target engagement was not measured on the profiled samples, so the weaker transcriptional response to TGX-221 cannot be read as evidence about the role of p110β.

#### Genotype of the model

NCI-H69 carries a PIK3CA in-frame deletion. A PIK3CA-altered line may have altered baseline reliance on p110α, so the findings cannot be generalised to PIK3CA-wild-type SCLC without empirical testing.

Two observations argue against a purely non-specific explanation. The response did not show the half-life dependence expected of a general transcriptional block, although that test is weak at 24 h. And the most strongly affected programme is one that PI3K–AKT–mTORC1 signalling is known to control. The most defensible reading is that the PIK-75 response mixes on-target PI3K inhibition with off-target effects in proportions these data cannot resolve.

Resolving them requires a selective p110α inhibitor such as alpelisib (42) or inavolisib (56), or p110α knockdown, profiled in the same way. The original study used siRNA against p110α for phenotypic assays, but not for the arrays.

### 4.3 The SREBP programme

mTORC1 promotes SREBP processing and, through it, sterol and fatty-acid synthesis (7, 8). The coordinated fall in SREBP targets, with little change in SREBF1 or SREBF2 mRNA, fits reduced SREBP activity rather than reduced SREBP expression. We measured transcripts only. Whether sterol or lipid synthesis itself falls, and whether cells then depend on exogenous lipid, was not tested. A direct prediction is that sensitivity to a selective p110α inhibitor increases in lipoprotein-depleted medium and is rescued by cholesterol.

### 4.4 Cell-cycle and homologous-repair genes

BRCA1 and several other homologous-repair genes decreased, but the effect was modest outside BRCA1, and core proliferation markers were unchanged. BRCA1 is cell-cycle regulated (57), and PIK-75 inhibits DNA-PK and CDK9. The data therefore cannot show that PI3Kα inhibition impairs homologous recombination. Reduced BRCA1/2 expression after PI3K inhibition has been reported in breast cancer models (58). Whether this occurs in SCLC would need a selective inhibitor and functional readouts such as RAD51 foci.

### 4.5 Dependency and clinical context

CRISPR screens show that SCLC lines depend more on PIK3CA than on PIK3CB, including lines without PIK3CA or PTEN mutations. Other lineages show the same hierarchy, so it is not a feature of SCLC. Lines with damaging PTEN mutations had the lowest PIK3CB scores, in line with the known reliance of PTEN-deficient tumours on p110β (5, 6).

The patient cohort gave no independent support. A score built from the PIK-75 response was nominally associated with survival, but random signatures of the same size performed similarly, a known property of expression signatures in outcome analyses (49). Tumours with PI3K-pathway mutations did not score differently. We therefore make no clinical claim.

PI3Kα inhibitors are approved for PIK3CA-mutant, hormone-receptor-positive breast cancer (9). In SCLC, activating PI3K-pathway alterations occur in a minority of tumours (2), and this study adds no evidence on which patients, if any, might benefit.

### 4.6 Limitations

#### One model

One PIK3CA-altered cell line, one 24-h time point and one dose per compound.

#### Compound selectivity

PIK-75 inhibits many kinases besides p110α. No selective inhibitor or genetic perturbation was profiled.

#### TGX-221 exposure

The concentration in the public record is ambiguous, and the response depended on two of three replicates. Target engagement was not measured on the profiled samples.

#### Design

Arrays were scanned in group order on one day, so scan order cannot be separated from treatment. Whether replicates were independent experiments is not recorded.

#### Normalisation

When most genes change, no normalisation recovers absolute changes per cell. Spike-in anchoring refers signal to the amount of material processed, not to cell number.

#### Probe-level ambiguity

For some genes the direction of change depends on the probe (for example MCL1 and FANCD2). We report these as unresolved (Table S7).

#### mRNA only

No protein, phosphorylation, metabolite or functional data were analysed.

#### Inferred activities

PROGENy and DoRothEA scores are indirect, and under broad repression most regulons score negative (64% of transcription factors here).

#### Validation

DepMap supports a general isoform hierarchy but did not screen NCI-H69. The cohort analysis was negative. No independent isoform-perturbation dataset from SCLC cells was analysed; the LINCS comparison used consensus signatures aggregated across cell lines and did not recover the signature of PIK-75 itself.

## 5. Declarations

### Ethics approval and consent to participate

This study re-analysed publicly available, de-identified data only. No new human participants, animals or samples were involved.

### Consent for publication

Not applicable.

### Data and code availability

All data are public: GEO series GSE40564; DepMap Public 24Q4; cBioPortal study sclc_ucologne_2015; LINCS L1000 consensus signatures obtained through Enrichr; and the consensus mRNA half-lives of Agarwal and Kelley. Analysis scripts, intermediate tables and session information are at https://github.com/Aref-Movaqar/sclc-pi3k-transcriptomics, archived at https://doi.org/10.5281/zenodo.23154076.

## Acknowledgements.

We thank the investigators who made GSE40564, DepMap and the UCologne SCLC cohort publicly available.

## References

1. Rudin CM, Brambilla E, Faivre-Finn C, Sage J. Small-cell lung cancer. Nat Rev Dis Primers. 2021;7(1):3.

2. George J, Lim JS, Jang SJ, Cun Y, Ozretić L, Kong G, et al. Comprehensive genomic profiles of small cell lung cancer. Nature. 2015;524(7563):47–53.

3. Rudin CM, Poirier JT, Byers LA, Dive C, Dowlati A, George J, et al. Molecular subtypes of small cell lung cancer: a synthesis of human and mouse model data. Nat Rev Cancer. 2019;19(5):289–97.

4. Knight ZA, Gonzalez B, Feldman ME, Zunder ER, Goldenberg DD, Williams O, et al. A pharmacological map of the PI3-K family defines a role for p110α in insulin signaling. Cell. 2006;125(4):733–47.

5. Jia S, Liu Z, Zhang S, Liu P, Zhang L, Lee SH, et al. Essential roles of PI(3)K-p110β in cell growth, metabolism and tumorigenesis. Nature. 2008;454(7205):776–9.

6. Wee S, Wiederschain D, Maira S-M, Loo A, Miller C, deBeaumont R, et al. PTEN-deficient cancers depend on PIK3CB. Proc Natl Acad Sci U S A. 2008;105(35):13057–62.

7. Porstmann T, Santos CR, Griffiths B, Cully M, Wu M, Leevers S, et al. SREBP activity is regulated by mTORC1 and contributes to Akt-dependent cell growth. Cell Metab. 2008;8(3):224–36.

8. Düvel K, Yecies JL, Menon S, Raman P, Lipovsky AI, Souza AL, et al. Activation of a metabolic gene regulatory network downstream of mTOR complex 1. Mol Cell. 2010;39(2):171–83.

9. André F, Ciruelos E, Rubovszky G, Campone M, Loibl S, Rugo HS, et al. Alpelisib for PIK3CA-Mutated, Hormone Receptor-Positive Advanced Breast Cancer. N Engl J Med. 2019;380(20):1929–40.

10. Wojtalla A, Fischer B, Kotelevets N, Mauri FA, Sobek J, Rehrauer H, et al. Targeting the phosphoinositide 3-kinase p110-α isoform impairs cell proliferation, survival, and tumor growth in small cell lung cancer. Clin Cancer Res. 2013;19(1):96–105.

11. Jackson SP, Schoenwaelder SM, Goncalves I, Nesbitt WS, Yap CL, Wright CE, et al. PI 3-kinase p110β: a new target for antithrombotic therapy. Nat Med. 2005;11(5):507–14.

12. Thomas D, Powell JA, Vergez F, Segal DH, Nguyen N-YN, Baker A, et al. Targeting acute myeloid leukemia by dual inhibition of PI3K signaling and Cdk9-mediated Mcl-1 transcription. Blood. 2013;122(5):738–48.

13. Lemke J, von Karstedt S, Abd El Hay M, Conti A, Arce F, Montinaro A, et al. Selective CDK9 inhibition overcomes TRAIL resistance by concomitant suppression of cFlip and Mcl-1. Cell Death Differ. 2014;21(3):491–502.

14. Shibata T, Kokubu A, Tsuta K, Hirohashi S. Oncogenic mutation of PIK3CA in small cell lung carcinoma: a potential therapeutic target pathway for chemotherapy-resistant lung cancer. Cancer Lett. 2009;283(2):203–11.

15. Barrett T, Wilhite SE, Ledoux P, Evangelista C, Kim IF, Tomashevsky M, et al. NCBI GEO: archive for functional genomics data sets—update. Nucleic Acids Res. 2013;41(Database issue):D991–D5.

16. Bairoch A. The Cellosaurus, a Cell-Line Knowledge Resource. J Biomol Tech. 2018;29(2):25–38.

17. Irizarry RA, Hobbs B, Collin F, Beazer-Barclay YD, Antonellis KJ, Scherf U, et al. Exploration, normalization, and summaries of high density oligonucleotide array probe level data. Biostatistics. 2003;4(2):249–64.

18. Gautier L, Cope L, Bolstad BM, Irizarry RA. affy—analysis of Affymetrix GeneChip data at the probe level. Bioinformatics. 2004;20(3):307–15.

19. Miller JA, Cai C, Langfelder P, Geschwind DH, Kurian SM, Salomon DR, et al. Strategies for aggregating gene expression data: the collapseRows R function. BMC Bioinformatics. 2011;12:322.

20. Ritchie ME, Phipson B, Wu D, Hu Y, Law CW, Shi W, et al. limma powers differential expression analyses for RNA-sequencing and microarray studies. Nucleic Acids Res. 2015;43(7):e47.

21. Smyth GK. Linear models and empirical Bayes methods for assessing differential expression in microarray experiments. Stat Appl Genet Mol Biol. 2004;3:Article3.

22. Phipson B, Lee S, Majewski IJ, Alexander WS, Smyth GK. Robust hyperparameter estimation protects against hypervariable genes and improves power to detect differential expression. Ann Appl Stat. 2016;10(2):946–63.

23. Benjamini Y, Hochberg Y. Controlling the false discovery rate: a practical and powerful approach to multiple testing. J R Stat Soc Series B Methodol. 1995;57(1):289–300.

24. McCarthy DJ, Smyth GK. Testing significance relative to a fold-change threshold is a TREAT. Bioinformatics. 2009;25(6):765–71.

25. Ritchie ME, Diyagama D, Neilson J, van Laar R, Dobrovic A, Holloway A, et al. Empirical array quality weights in the analysis of microarray data. BMC Bioinformatics. 2006;7:261.

26. Wu D, Lim E, Vaillant F, Asselin-Labat M-L, Visvader JE, Smyth GK. ROAST: rotation gene set tests for complex microarray experiments. Bioinformatics. 2010;26(17):2176–82.

27. Wu D, Smyth GK. Camera: a competitive gene set test accounting for inter-gene correlation. Nucleic Acids Res. 2012;40(17):e133.

28. Korotkevich G, Sukhov V, Budin N, Shpak B, Artyomov MN, Sergushichev A. Fast gene set enrichment analysis. bioRxiv. 2021:060012.

29. Subramanian A, Tamayo P, Mootha VK, Mukherjee S, Ebert BL, Gillette MA, et al. Gene set enrichment analysis: a knowledge-based approach for interpreting genome-wide expression profiles. Proc Natl Acad Sci U S A. 2005;102(43):15545–50.

30. Liberzon A, Birger C, Thorvaldsdóttir H, Ghandi M, Mesirov JP, Tamayo P. The Molecular Signatures Database (MSigDB) hallmark gene set collection. Cell Syst. 2015;1(6):417–25.

31. Milacic M, Beavers D, Conley P, Gong C, Gillespie M, Griss J, et al. The Reactome Pathway Knowledgebase 2024. Nucleic Acids Res. 2024;52(D1):D672–D8.

32. Schubert M, Klinger B, Klünemann M, Sieber A, Uhlitz F, Sauer S, et al. Perturbation-response genes reveal signaling footprints in cancer gene expression. Nat Commun. 2018;9(1):20.

33. Badia-i-Mompel P, Vélez Santiago J, Braunger J, Geiss C, Dimitrov D, Müller-Dott S, et al. decoupleR: ensemble of computational methods to infer biological activities from omics data. Bioinform Adv. 2022;2(1):vbac016.

34. Garcia-Alonso L, Holland CH, Ibrahim MM, Turei D, Saez-Rodriguez J. Benchmark and integration of resources for the estimation of human transcription factor activities. Genome Res. 2019;29(8):1363–75.

35. Agarwal V, Kelley DR. The genetic and biochemical determinants of mRNA degradation rates in mammals. Genome Biol. 2022;23(1):245.

36. Subramanian A, Narayan R, Corsello SM, Peck DD, Natoli TE, Lu X, et al. A Next Generation Connectivity Map: L1000 Platform and the First 1,000,000 Profiles. Cell. 2017;171(6):1437–52.e17.

37. Evangelista JE, Clarke DJB, Xie Z, Lachmann A, Jeon M, Chen K, et al. SigCom LINCS: data and metadata search engine for a million gene expression signatures. Nucleic Acids Res. 2022;50(W1):W697–W709.

38. Xie Z, Bailey A, Kuleshov MV, Clarke DJB, Evangelista JE, Jenkins SL, et al. Gene Set Knowledge Discovery with Enrichr. Curr Protoc. 2021;1(3):e90.

39. Parry D, Guzi T, Shanahan F, Davis N, Prabhavalkar D, Wiswell D, et al. Dinaciclib (SCH 727965), a novel and potent cyclin-dependent kinase inhibitor. Mol Cancer Ther. 2010;9(8):2344–53.

40. Thoreen CC, Kang SA, Chang JW, Liu Q, Zhang J, Gao Y, et al. An ATP-competitive mammalian target of rapamycin inhibitor reveals rapamycin-resistant functions of mTORC1. J Biol Chem. 2009;284(12):8023–32.

41. Liu Q, Wang J, Kang SA, Thoreen CC, Hur W, Ahmed T, et al. Discovery of 9-(6-aminopyridin-3-yl)-1-(3-(trifluoromethyl)phenyl)benzo[h][1,6]naphthyridin-2(1H)-one (Torin2) as a potent, selective, and orally available mammalian target of rapamycin (mTOR) inhibitor for treatment of cancer. J Med Chem. 2011;54(5):1473–80.

42. Fritsch C, Huang A, Chatenay-Rivauday C, Schnell C, Reddy A, Liu M, et al. Characterization of the novel and specific PI3Kα inhibitor NVP-BYL719 and development of the patient stratification strategy for clinical trials. Mol Cancer Ther. 2014;13(5):1117–29.

43. Dempster JM, Boyle I, Vazquez F, Root DE, Boehm JS, Hahn WC, et al. Chronos: a cell population dynamics model of CRISPR experiments that improves inference of gene fitness effects. Genome Biol. 2021;22(1):343.

44. Tsherniak A, Vazquez F, Montgomery PG, Weir BA, Kryukov G, Cowley GS, et al. Defining a Cancer Dependency Map. Cell. 2017;170(3):564–76.e16.

45. DepMap B. DepMap 24Q4 Public. Figshare+; 2024.

46. Cerami E, Gao J, Dogrusoz U, Gross BE, Sumer SO, Aksoy BA, et al. The cBio cancer genomics portal: an open platform for exploring multidimensional cancer genomics data. Cancer Discov. 2012;2(5):401–4.

47. Gao J, Aksoy BA, Dogrusoz U, Dresdner G, Gross B, Sumer SO, et al. Integrative analysis of complex cancer genomics and clinical profiles using the cBioPortal. Sci Signal. 2013;6(269):pl1.

48. Zhang W, Girard L, Zhang Y-A, Haruki T, Papari-Zareei M, Stastny V, et al. Small cell lung cancer tumors and preclinical models display heterogeneity of neuroendocrine phenotypes. Transl Lung Cancer Res. 2018;7(1):32–49.

49. Venet D, Dumont JE, Detours V. Most random gene expression signatures are significantly associated with breast cancer outcome. PLoS Comput Biol. 2011;7(10):e1002240.

50. Team RC. R: a language and environment for statistical computing. 4.4.3 ed. Vienna, Austria: R Foundation for Statistical Computing; 2025.

51. Davis S, Meltzer PS. GEOquery: a bridge between the Gene Expression Omnibus (GEO) and BioConductor. Bioinformatics. 2007;23(14):1846–7.

52. Therneau TM. A package for survival analysis in R. R package version 3.8 ed2024.

53. Chandarlapaty S, Sawai A, Scaltriti M, Rodrik-Outmezguine V, Grbovic-Huezo O, Serra V, et al. AKT inhibition relieves feedback suppression of receptor tyrosine kinase expression and activity. Cancer Cell. 2011;19(1):58–71.

54. Serra V, Scaltriti M, Prudkin L, Eichhorn PJA, Ibrahim YH, Chandarlapaty S, et al. PI3K inhibition results in enhanced HER signaling and acquired ERK dependency in HER2-overexpressing breast cancer. Oncogene. 2011;30(22):2547–57.

55. Brunet A, Bonni A, Zigmond MJ, Lin MZ, Juo P, Hu LS, et al. Akt promotes cell survival by phosphorylating and inhibiting a Forkhead transcription factor. Cell. 1999;96(6):857–68.

56. Turner NC, Im S-A, Saura C, Juric D, Loibl S, Kalinsky K, et al. Inavolisib-Based Therapy in PIK3CA-Mutated Advanced Breast Cancer. N Engl J Med. 2024;391(17):1584–96.

57. Gudas JM, Li T, Nguyen H, Jensen D, Rauscher FJ, Cowan KH. Cell cycle regulation of BRCA1 messenger RNA in human breast epithelial cells. Cell Growth Differ. 1996;7(6):717–23.

58. Ibrahim YH, García-García C, Serra V, He L, Torres-Lockhart K, Prat A, et al. PI3K inhibition impairs BRCA1/2 expression and sensitizes BRCA-proficient triple-negative breast cancer to PARP inhibition. Cancer Discov. 2012;2(11):1036–47.

